# Breaking down defenses: quantitative analysis of malaria infection dynamics reveals distinct immune defense strategies

**DOI:** 10.1101/648428

**Authors:** Nina Wale, Matthew J. Jones, Derek G. Sim, Andrew F. Read, Aaron A. King

**Author notes:** Author contributions: NW and AAK designed the research, performed the analysis, and wrote the manuscript. MJJ and DGS performed the experiment and contributed to its design. AFR, MJJ, and DGS provided comments.

## Abstract

Hosts defend themselves against pathogens by mounting an immune response. Fully understanding the immune response as a driver of host disease and pathogen evolution requires a quantitative account of its impact on parasite population dynamics. Here, we use a data-driven modeling approach to quantify the birth and death processes underlying the dynamics of infections of the rodent malaria parasite, *Plasmodium chabaudi*, and the red blood cells (RBCs) it targets. We decompose the immune response into three components, each with a distinct effect on parasite and RBC vital rates, and quantify the relative contribution of each component to host disease and parasite density. Our analysis suggests that these components are deployed in a coordinated fashion to realize distinct resource-directed defense strategies that complement the killing of parasitized cells. Early in the infection, the host deploys a strategy reminiscent of siege and scorched-earth tactics, in which it both restricts the supply of RBCs and destroys them. Late in the infection, a ‘juvenilization’ strategy, in which turnover of RBCs is accelerated, allows the host to recover from anemia while holding parasite proliferation at bay. By quantifying the impact of immunity on both parasite fitness and host disease, we reveal that phenomena often interpreted as immunopathology may in fact be beneficial to the host. Finally, we show that, across mice, the components of the host response are consistently related to each other, even when infections take qualitatively different trajectories. This suggests the existence of simple rules that govern the immune system’s deployment.

---

When a pathogen infects a host, the host mounts a response to contain the pathogen and protect itself from harm. This response shapes the within-host ecological milieu in which pathogens proliferate and evolve and, in some cases, can directly contribute to host disease, a phenomenon known as immunopathology (Hochberg, 2018; Graham et al., 2005). To fully understand host-parasite interactions, infection pathology, and host-parasite coevolution, we must therefore quantify the nature and dynamics of the host response to infection (Hedrick, 2017).

Yet quantifying the impact of immunity on host and parasite fitness remains a challenge. Mechanistic approaches to the study of immunity, that characterize the host-cell populations involved and quantify the mechanisms responsible for changes in their abundance, distribution, and activity, have had many successes, including some that have informed vaccines and immunotherapeutic treatments. However, even equipped with an understanding of a cell type’s function, it can be difficult to quantify that cell type’s impact on parasite abundance or host symptoms as a function of its density (Zinkernagel, 2007; Hochberg, 2018). Here, we take a complementary approach that explicitly focusses on the net effects of the host response on births and deaths of host and parasite cells which, in turn, directly affect host health and parasite fitness. Specifically, we decompose the host response to the mouse malaria parasite, *Plasmodium chabaudi*, into qualitatively distinct components that variously kill parasites or control their access to resources. We use a data-driven approach to extract the time-course of each component from experimental data and then compute the impacts of each on both host symptoms (i.e., anemia) and parasite proliferation. In so doing, we paint a quantitative picture of the host response as both a driver of disease and a selective pressure on parasites. We examine how the trajectories of the distinct components relate to parasite burden and to each other and formulate hypotheses as to how and to what end they are deployed. By thus focusing on the effects of the host response as distinct from effectors that mediate it, we elucidate the strategies a host employs to control an infection, as distinct from the weaponry it uses to achieve them.

## Results

### Decomposition of the host response

We developed a simple, semi-mechanistic approach to infer the nature and shape of the host response to malaria infections from experimental data (*Materials & Methods*). We take advantage of *P. chabaudi*’s synchronous, daily life-cycle: Every 24 h, these parasites invade, repro-duce, and burst out of red blood cells (RBCs), which are destroyed in the process (Fig. 1A). With measures of the densities of parasites and RBCs, together with the rate at which immature RBCs (reticulocytes) are supplied to the bloodstream (Fig. 1B), and supposing that parasite reproduction is limited by RBC availability alone, one can forecast how many parasitized and unparasitized RBCs should be present a day later. Observed deviations from this forecast can thence be attributed either to a host response targeted at parasitized cells (‘targeted killing’) or to one that removes RBCs irrespective of their infection status (‘indiscriminate killing’; Fig. 1C and *Materials & Methods*). Repeatedly applying this process to time series data collected from experimental infections, we deduce the trajectories of each of three functionally distinct, time-varying components (Fig. 1D): targeted killing, indiscriminate killing, and RBC supply restriction.

**Figure 1.**
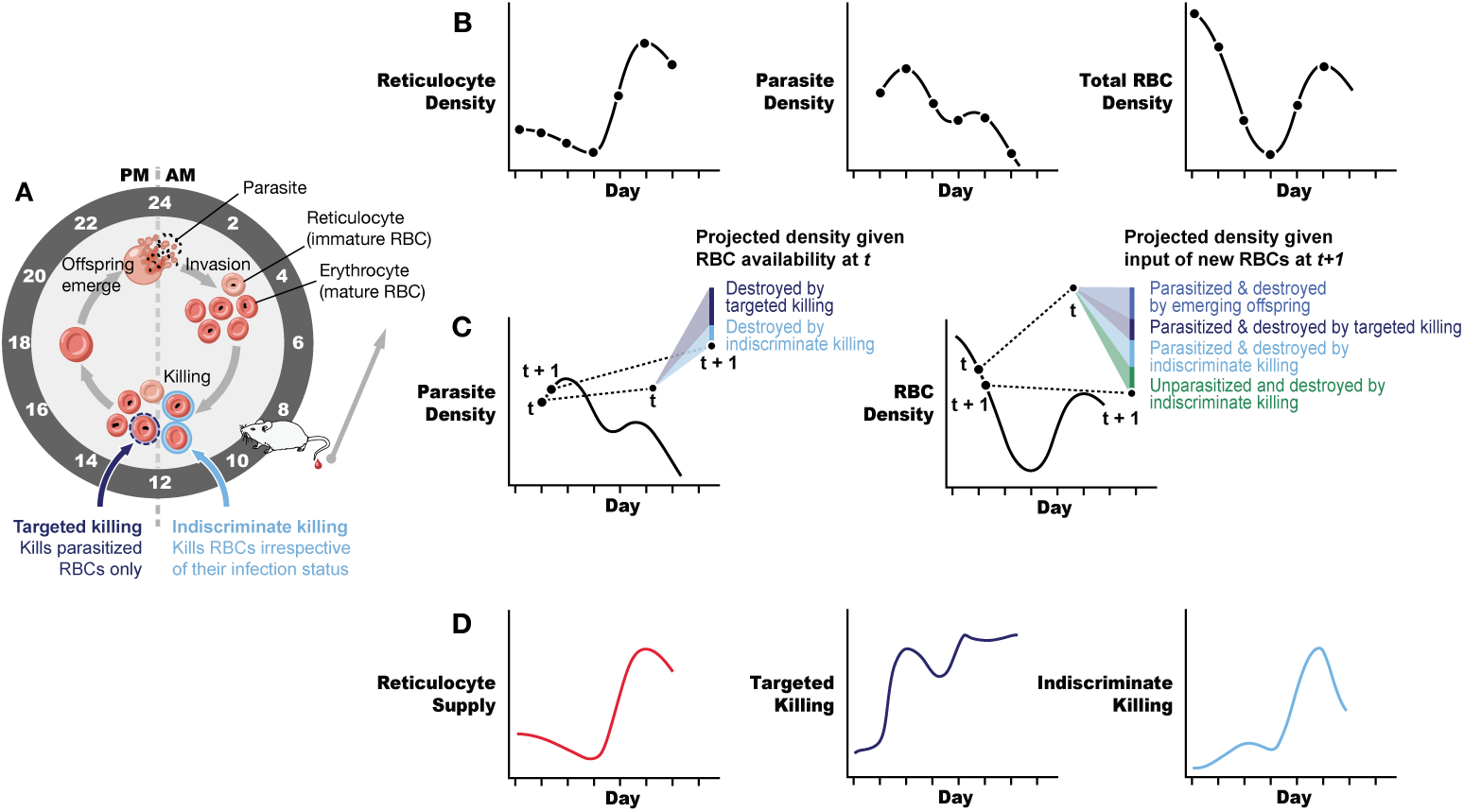
Decomposing the host immune response. A) The daily cycle of malaria parasites in the blood stage of infection. Each day, parasite offspring (merozoites) burst from red blood cells (RBCs) and then attempt to invade new RBCs, wherein they mature and reproduce. The host immune response modulates infection by, among other things, killing RBCs and adjusting their supply. Samples are taken from experimentally infected mice each morning before parasites have reproduced and are processed to produce B) Time series of reticulocyte (i.e., immature RBC), parasite, and total (mature+immature) RBC densities. C) Eq. 1 shows how these three data streams can be transformed into three synthetic variables (components of the immune response), which describe the impacts of the immune response on parasite proliferation and host health (anemia). In particular, RBCs can be killed by a response targeted at parasitized cells or by a response that kills RBCs irrespective of their infection status. Moreover, the host can modulate the supply of reticulocytes. The model works by projecting the density of RBC and parasites at the next time step (*t+1*) in the absence of any killing, given the abundance of RBCs and parasites at time *t*, the supply of RBCs at time *t+1*. The deficit between these projections and the data can is then partitioned among the indiscriminate and targeted killing components. D) This procedure yields, for each infected mouse, time series of the three immune response components. By comparing these trajectories across mice, we identify robust patterns which can be interpreted in terms of host defense strategy.

We applied this approach to data (densities of reticulocytes, total RBCs and parasites) collected daily from 12 experimentally-infected and 3 uninfected mice of the same genetic background (Fig. S-1). To generate variability in the infection dynamics among the mice, they were split into four groups, each of which was supplied with a different concentration of a nutrient (para-aminonenzoic acid, herafeter pABA) that alters the growth rate and dynamics of parasite populations but is not needed by the host (Wale et al.,Fenton et al., 1950). For brevity, in the main text, we show and analyze data from the same three mice, each of which was given a different concentration of the nutrient; the complete data and analysis of all mice can be found in the *Supplementary Information*.

Mice exhibited markedly different infection dynamics. As expected, the initial growth rate of infections increased with the concentration of pABA that they were administered (*F*_3,29_ = 5.4, *p <* 0.01). Furthermore, while some mice experienced severe anemia, i.e., pathologically low numbers of RBCs (e.g., Fig. 2A, mouse 1), or displayed a ‘shoulder’ in the post-peak phase of the infection (e.g., Fig. 2A, mice 1–2), others did not (e.g. Fig. 2A, mouse 3).

**Figure 2.**
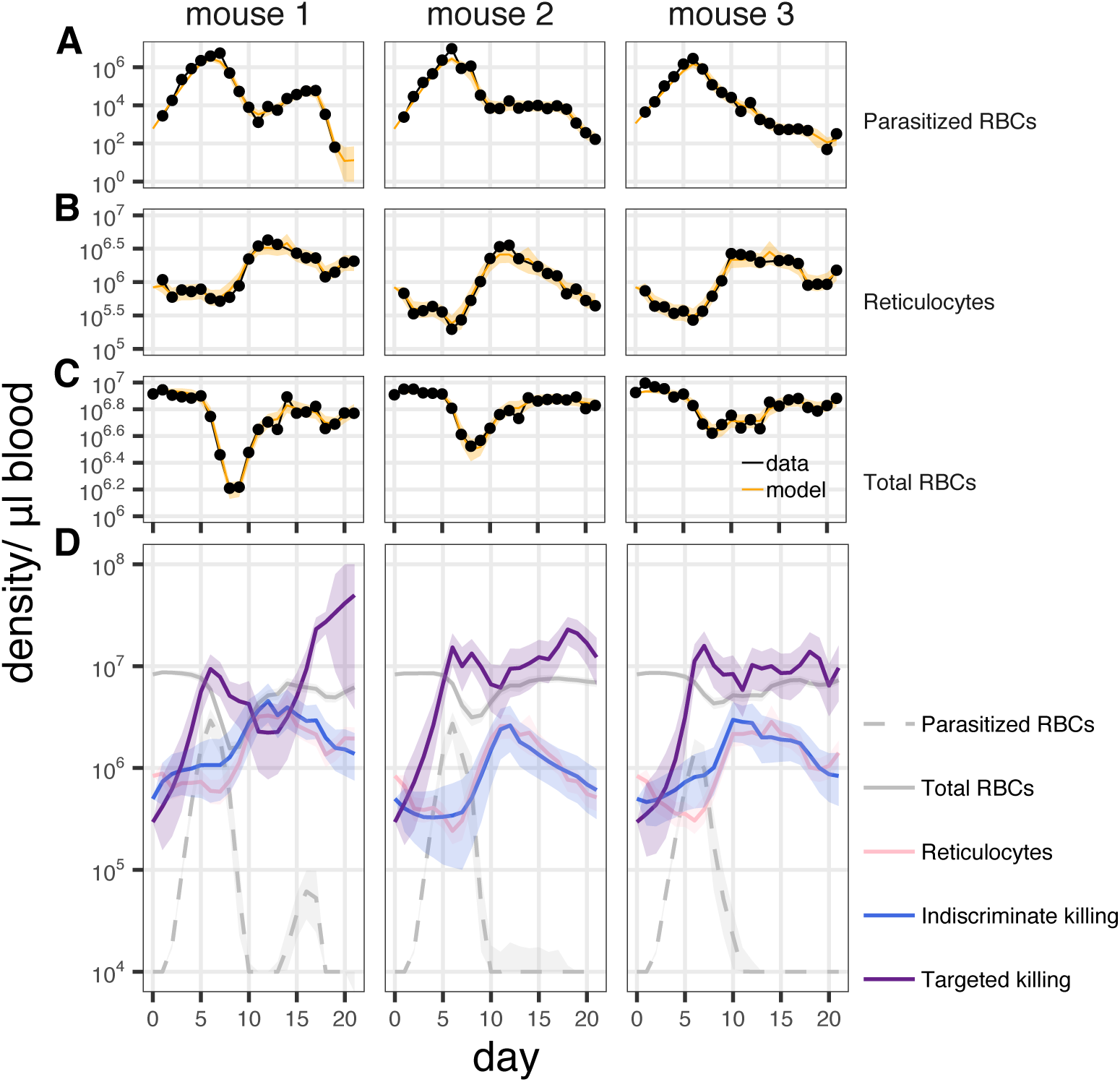
The model accurately captures the data and yields the dynamics of three qualitatively distinct host responses. Measured (black) and fitted (orange) densities of A) parasitized RBCs, B) reticulocytes and C) total RBCs in three mice which were (left to right) fed a 0.005% concentration, 0.0005% concentration and a 0% solution of pABA, a nutrient that stimulates parasite growth rate. Data and fitted model trajectories for all mice are shown in Fig. S-1 & S-2. D) Estimated trajectories of three distinct host responses that target parasitized cells only (”targeted killing”, purple), RBCs irrespective of their infection status (indiscriminate killing, blue) and that resupply reticulocytes (pink). Note that these responses are of the same order of magnitude as the RBCs. Plotted are the mean (solid line) and 90% confidence interval (ribbon) on the smoothed estimate of the model trajectories.

The model is sufficiently flexible that the dynamics of infection of all mice are well captured (Fig. 2A-C). Despite the variation in infection dynamics, the trajectories of the three response components are qualitatively similar across mice (Fig. 2D). These functions are scaled to be commensurate with RBC densities so that their magnitudes relative to RBC density determine the fate of RBCs and parasites (cf. Eqs. 1). In the pre-peak phase (*∼* 0–7d), the targeted response increases rapidly and peaks in concert with parasite density, while the indiscriminate response shows no discernible trend. Meanwhile, the supply of reticulocytes goes unchanged or even, in some cases, falls. Following the peak in parasite density, the targeted response falls as the indiscriminate response and reticulocyte supply increase. Strikingly, the trajectories of the latter two components move in tandem during this period (Fig. 2D). Once RBC numbers have recovered (*∼* day 15), the indiscriminate and supply responses decrease in magnitude and the targeted response increases once again.

### Parsing the effect of the host response on pathology and parasite fitness

To better understand the impact of each of the response components on parasite fitness and host disease, and hence to identify putative host defense strategies, we quantify the effect of each response component on parasite destruction and on anemia (Fig. 3).

**Figure 3.**
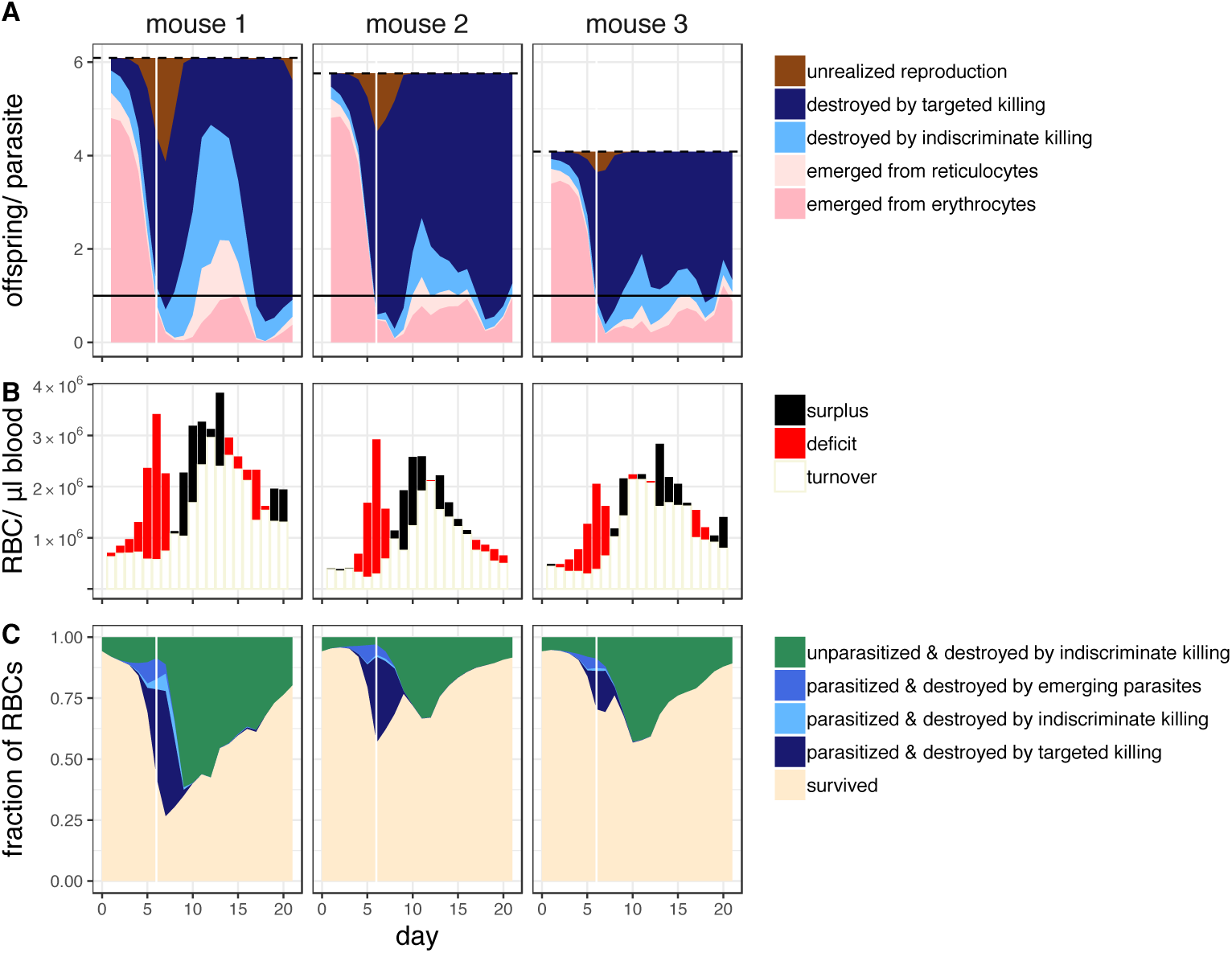
Contribution of the different components of the host response to parasite fitness and host disease. A) Contribution of the host to the (suppression of) parasite reproduction in three different mice (left to right), each of which was fed a different concentration of a nutrient that stimulates parasite growth. Dashed line indicates the maximum number of offspring that could be produced per parasitized cell in conditions of unlimited RBC availability; fill area indicates the estimated number of offspring that successfully emerged (pinks), were destroyed by each of the killing responses (blues) or were not produced due to the limited availability of RBCs (brown). Black horizontal line indicates one offspring/parasite: when the blue area descends below this line the parasite population is decreasing in size. B) Uncolored part of the bar indicates the number of the previous day’s losses that were compensated for by the present day’s supply of reticulocytes (turnover); the colored section indicates the extent to which the present day’s supply of reticulocytes exceeded yesterday’s losses (‘surplus’, black) or failed to compensate for them (‘deficit’, red). C) The relative contribution of parasites and each of the host responses to the fate of RBCs through time. Note that parasites can contribute to RBC destruction directly, by emergence, or indirectly via the targeted killing response. White vertical line in A & C indicates the time of peak parasite density. Shown are the decompositions for the same three mice used in all other figures. Equivalent plots for the nine other mice analyzed can be found in (Fig. S-3).

At the infection’s outset, parasite populations grow at near their maximum rate and the majority of parasite offspring emerge from mature RBCs (erythrocytes). Very rapidly, however, parasites are unable to realize their reproductive potential due to the combined impacts of the targeted response and RBC limitation. At its maximum, RBC limitation accounts for, on average, a 22% (interdecile range, 12–33%) reduction in parasite reproductive potential (Fig. 3A).

RBC limitation results from RBC destruction by the parasite and the host, as well as the restriction of RBC supply. Together, these three forces induce anemia. Notably, only a small fraction of RBC losses are attributable to parasites bursting from RBCs (Fig. 3C). Even when the contribution of parasite emergence to RBC destruction is at its height, it is never responsible for more than half of the RBC destruction (maximum (across time) proportion of losses attributable to parasite emergence, mean (among mice) 33%, interdecile range 18%–43%). Rather, most RBC losses are due to the indiscriminate response, which removes both parasitized and unparasitized cells. At the beginning of the infection, almost all of the losses of RBCs can be attributed to the removal of uninfected cells by the indiscriminate response (mean, 99.5%, interdecile range 99–100%) and indeed there are only a few days, around the time of peak parasite density, when the host kills more parasitized cells than uninfected cells. On these days, 4 parasitized cells are removed on average for every 1 unparasitized cell (interdecile range 1.3–9). In addition, the restriction of reticulocyte supply exacerbates anemia, as each day’s supply of new RBCs fails to compensate for the previous day’s losses (Fig. 3B). Strikingly, despite incurring huge losses in RBCs, infected mice do not increase the supply of reticulocytes any more than uninfected mice in the first 8 days of infection (Fig. 2, *total reticulocyte count, days* 0–8, *F*_1,13_ = 3.2, *p* = 0.1). Metaphorically speaking, the restriction of RBC supply and the killing of uninfected RBCs represent “siege” and “scorched earth” strategies, respectively, which combine to limit the resources available to the parasite and make the host sick. These strategies supplement the “slaughter” of infected cells by the targeted response to bring the parasite population under control.

The post-peak phase of infection, in which parasite numbers decline and mice recover from anemia, is characterized by an increase in the magnitude of the indiscriminate response. At its peak, this response component is responsible for 53% (interdecile range 38–70%) of parasite destruction on average (Fig. 3A), as well as the destruction of many unparasitized RBCs.

Despite this post-peak increase in indiscriminate killing, the mouse recovers from anemia. This is due to the concomitant increase in the supply of reticulocytes, which more than compensates for the destruction of RBCs (Fig. 3B). In contrast to the pre-peak phase, here the host controls parasite population growth not by making RBCs scarce but by altering their demographic profile. Specifically, because the indiscriminate response increases in tandem with reticulocyte supply, the combined effect is to increase the turnover rate of RBCs, markedly reducing their average age. Thus, although RBCs become more abundant, parasites that successfully invade an RBC still have a low probability of reproducing, since the RBC that they have invaded is likely to be cleared before they do so. Strategically speaking, this “juvenilization” strategy allows the host to recover from anemia while restraining parasite growth.

In the final phase (days 15–21), both the indiscriminate killing and supply responses return to roughly their initial levels. The targeted response, by contrast, remains at heightened levels and, in some cases, increases.

### Coordination of the host response

Though infection dynamics vary among mice, the components of the host response follow stereotypical trajectories (Fig. 2D). This suggests that the components may be deployed in a coordinated manner. To investigate this possibility, we plotted the trajectories of each response against one another and against parasite density. This analysis reveals that the infection rolls out in three phases, separated by two landmarks (denoted **a** and **b** in Fig. 4).

**Figure 4.**
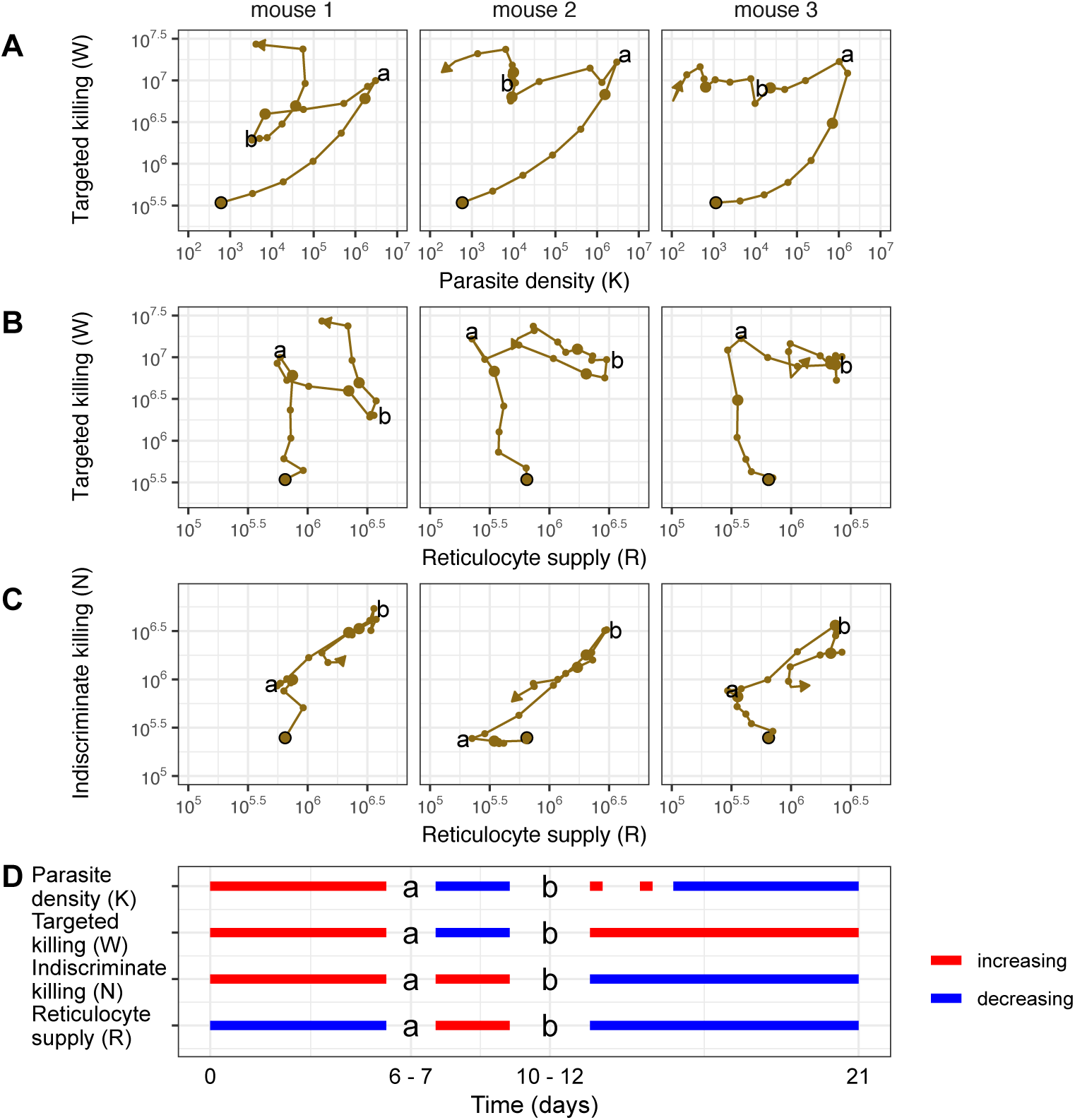
The components of the immune response are similarly deployed, even among mice displaying different infections dynamics. (A-C) The relationships between the three components of the host response. Large, black-lined dot indicates the starting point of the infection. Each subsequent day is marked with a dot with large dots indicating days 5, 10, 15, 20. Landmarks **a** and **b** identify transitions between regimens of deployment of the components that occur, as summarized in D (NB: the timing of these landmarks are consistent across all mice in the analysis, with the exception of two mice that received fewer parasites than intended, see Figs. S-1 & S-4). In D, dashed red line indicates that parasite density increases in some but not all mice during the period indicated. Densities of parasites and RBCs are in units per microliter of blood. These trajectories are from the same three mice whose infections are displayed in Figs. 2 & 3, each of which received a different concentration of pABA, a nutrient that alters parasite growth rate.

In the first phase, the targeted response grows with the parasite population, as the reticulocyte supply falls and the indiscriminate response slowly grows. At landmark **a**, the parasite population and targeted killing response stop growing and begin to decrease. Notably, there is no time lag between the fall in parasite density and the fall in the targeting killing response (Fig. 4A). Landmark **a** also marks the point where the magnitude of the reticulocyte supply and indiscriminate killing responses become tightly correlated, growing and waning with each other for the duration of the infection (Fig. 4C). Landmark **b** marks the beginning of a third phase, where the indiscriminate response reaches its peak and begins to fall, along with the supply of reticulocytes. The growth rate of the parasite population and targeted killing response become decoupled and their trajectories more idiosyncratic, in this phase. While the targeted response tends to grow during this latter phase, only in some some mice does parasite density stop falling or grow again, so as to form a ‘shoulder’ in the parasite density curve, before finally falling to undetectable levels.

## Discussion

Understanding the character and dynamics of the host response to infection is essential if we are to apportion responsibility for pathology to parasite and host, elucidate the role of the immune response in shaping parasite traits, and, ultimately, predict the dynamics of infections. Here, we used a simple accounting scheme to decompose the host response into three functionally distinct components and to infer their respective dynamics. This decomposition allows us to formulate hypotheses regarding the strategic significance of aspects of the immune response.

Our analysis suggests that host responses to infection that have been interpreted as immunopathology may in fact serve as effective defense strategies. The restriction of the supply of reticulocytes to the bloodstream at the outset of malaria infections, the accelerated clearance of erythrocytes toward the end of infections and the destruction of uninfected erythrocytes are well-known features of malaria infections in primates and rodents (e.g. (Chang and Stevenson, 2004; Salmon et al., 1997; Jakeman et al., 1999; Looareesuwan et al., 1987, 1991)). Each have, at times, been interpreted as pathological. For example, restricted erythropoeisis has been dubbed “inappropriate” or “ineffective” erythropoiesis in the literature (Chang and Stevenson, 2004; Deroost et al., 2016). Our analysis provides new evidence that, while these phenomena do contribute to the anemia of malaria infections (Fig. 3BC), they also contribute importantly to the control of parasite populations. In particular, early in an infection, the restriction of reticulocyte supply and the destruction of uninfected cells shrink the reproductive potential of parasites; in the later phase, increased RBC clearance, in conjunction with augmented erythropoiesis, accelerates blood compartment turnover, thereby reducing parasite survival while allowing the host to recover. Our analysis thus illustrates the general principle that, in evaluating whether a given component of the host response is pathological or not, one must consider it in the context of other, simultaneously mounted, responses and its effects on parasite fitness as well as host disease. Indeed some aspects of illness may, when considered in this way, prove to be adaptive (Ewald, 1994). The relative contributions of resource limitation (“bottom-up control”) and direct killing (“top-down control”) to the dynamics of malaria infections has been hotly debated, with several authors suggesting that RBC limitation alone is sufficient to explain infection dynamics in the acute phase (Antia et al., 2008; Mideo et al., 2008; Metcalf et al., 2011). While our model is flexible enough to encompass such an explanation, we find that the model most consistent with the data is one in which the host slows parasite population growth in the acute phase by a combination of resource limitation and direct removal of parasitized cells. We suspect that this discrepancy with previous findings arises from our use of explicit measurements of reticulocyte supply, where previous efforts relied on model assumptions. Similarly, the absence of data on reticulocyte abundance from previous studies likely explains why resource turnover has not been previously highlighted as an important mechanism of regulating malaria infections. To further examine the role of resource limitation in limiting parasite growth, we suggest that experimental manipulation of RBC densities be performed. For example, our work predicts that the transfusion of RBCs would not significantly boost parasite densities, except during the short period where resource limitation significantly impacts parasite fitness (i.e., within the 4–5 day window surrounding the peak in parasite density; Fig. 3A).

Though there is variation in the relative contribution of the different components of the host response to infection dynamics, the relationships between these components are consistent across mice, suggesting that simple, predictive models of infection dynamics might be built based on this work. Interestingly, we uncover patterns of co-regulation that contravene the standard assumptions of many models of within-host infection dynamics. Specifically, mechanistic models of immunity have commonly described the immune response as a predator-prey interaction, whereby the immune response (the “predator”) grows as a function of the density of the infected cells (the “prey”) (Bell, 1973; Wodarz, 2006; Fenton and Perkins, 2010). Our analysis (Fig. 4) shows that the waxing and waning of the targeted killing response is not a function of parasite abundance alone. A broader class of models may thus be required to accurately capture the within-host dynamics of infections.

A complete understanding of the immune response requires that we not only map its effects on parasite and host cells, as we do here, but also understand the mechanisms that mediate those effects. Intriguingly, not only does our model describe phenomena common in the malaria literature (see above) but the patterns of co-regulation echo findings in the wider, immunological literature. For example, the antagonism between the parasite killing response/parasite density and the supply response (Fig. 4D) is reminiscent of the antagonism of erythropoiesis by TNFα and hemozoin, which halt the production of reticulocytes (Clark and Chaudhri, 1988; Casals-Pascual et al., 2006).

The identification of the mechanisms underlying observed patterns is unlikely to be routinely straight-forward, however. We expect that, in general, multiple cell types and signaling molecules will mediate the responses we describe. For example, there are several mechanisms that might be responsible for the increased removal of uninfected cells during malaria infections (Salmon et al., 1997; Safeukui et al., 2015; Fernandez-Arias et al., 2016). Experimental manipulation of infection dynamics, in combination with measurement of a panel of putative mediators (as per the procedures used in systems immunology (Davis et al., 2017)), has the potential to tease apart which of the array of cell types are the most sensitive predictors of each of these functional components of the host response. Population-level approaches, such as ours, that focus on the quantitative effects of the immune response are an invaluable complement to cellular-level approaches focused on immune mechanisms. By combining these approaches, we stand to gain a holistic understanding of infection and immunity.

## Materials and Methods

### Hosts and parasites

Hosts were 15 6-8 week old C57BL/6J females. Twelve mice were infected intraperitoneally with 10^6^ *Plasmodium chabaudi* parasites of the pyrimethamine-resistant AS_124_ strain; 3 mice were left uninfected but received a sham injection of DMSO to control for effects of receiving an injection. To create variation in infection dynamics, three mice were assigned to each of four treatments that received a 0.05%, 0.005%, 0.0005%, or 0% solution of para-aminobenzoic acid (pABA) as drinking water, from a week before parasites were inoculated. Uninfected mice received a 0% pABA solution as drinking water.

### Infection monitoring

Infections were monitored daily from the day of inoculation (day 0) to day 21 post-inoculation (PI) and then every other day until day 29. 14 µl of blood was taken from the tail: 5 µl for the assessment of parasite density via quantitative PCR (qPCR) and 2 µl for the quantitation of total RBC density via Coulter counter (Beckman Coulter), as previously described Wale et al. (2017). A further 2 µl was used in the quantitation of young RBC (see below) and the remainder for other assays, not described herein. Since all of the mice had recovered from anemia (save one that died on day 9) and the majority had cleared the infection by day 21, we restrict our discussion to the first 21 days of the infection.

The proportion of RBCs that were reticulocytes (young RBCs) and erythrocytes (mature RBCs) was assayed using flow cytometry. Reticulocytes possess the CD71 (Transferrin Receptor 1), which is lost upon their maturation into erythrocytes (Pan et al., 1983; Liu et al., 2010); all cells in the erythroid lineage express the TER119 antigen Kina et al. (2000). Staining blood with TER-119 and CD71 antibodies labelled with distinct fluorophores was thus used to isolate the RBC portion of the blood and measure the relative abundance of reticulocytes and erythrocytes Koulnis et al. (2011). Briefly, 2 µl blood was collected in 48 µl running buffer (PBS with 2 mM EDTA and 2% FBS). Cells were mixed with 50 µl of each of three different solutions: FITC anti-mouse CD71 (Biolegend 113805), PE anti-mouse TER-119 (Biolegend 116207), APC anti-mouse CD41 (Biolegend 133914), for final concentrations of 0.005, 0.0025, and 0.0025 µg/µl, respectively. Samples were vortexed and incubated for an hour at 4C in the dark, washed with 1 ml of running buffer and centrifuged at 2000 rpm for 5 min. The supernatant was then decanted and the pellet resuspended to a final concentration of 10^7^ cells/µl. Samples were analyzed on a FlowSight Imaging Flow Cytometer (Amnis) according to the method and specifications described in the supplementary text. To obtain densities of reticulocytes and erythrocytes, the proportions obtained from the above analyses were multiplied by the total RBC density.

### Model structure

To decompose the host response to infection, we formulated a model of the interaction among RBCs, parasites, and three host response components. The model tracks the concentrations of erythrocytes, *E*_*t*_, reticulocytes, *R*_*t*_, and merozoites (parasite offspring), *M*_*t*_, as functions of time *t*. Given these quantities, together with the values on day *t* of the targeted and indiscriminate killing responses (*W*_*t*_, *N*_*t*_, respectively), the module postulates that

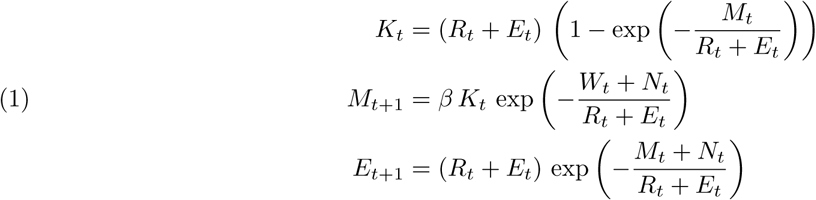

where *K*_*t*_ is the concentration of parasitized RBCs. Following empirical work (Jarra and Brown, 1989; Yap and Stevenson, 1994), we assume that all RBCs are equally susceptible to the parasite. According to Eqs. 1, each of the *K*_*t*_ parasitized cells contributes *β* merozoites to the population of day *t* + 1, if it is not first removed by one of the two killing responses (*N*_*t*_, *W*_*t*_). Note that the magnitudes of the killing responses are expressed in units of cell density. The third host response component is the supply of new reticulocytes into the bloodstream, *R*. We assume that reticulocytes mature into erythrocytes in precisely one day (Ney, 2011). The form of Eqs. 1 arises as the expectation of a stochastic urn process whereby each red blood cells faces a chance of becoming parasitized or destroyed by the host response in proportion to the relative abundances of *W*_*t*_, *N*_*t*_, and *M*_*t*_. As such, Eqs. 1 are deterministic, conditional on *N*_*t*_, *W*_*t*_, and *R*_*t*_. Stochasticity enters the model via the assumption that these variables are Gaussian Markov random fields (GMRF), with respective standard deviations *σ*_*N*_, *σ*_*W*_, *σ*_*R*_. Finally, the variables *R, E*, and *K* are related to the data via an explicit model of measurement errors. Specifically, measurements of parasite, reticulocyte, and total RBC densities on day *t* are assumed to be log-normally distributed around their true values (*K*_*t*_, *R*_*t*_, and *E*_*t*_ + *R*_*t*_, respectively).

### Model fitting and smoothing

The model has the form of a partially observed Markov process (King et al., 2016), for which efficient inference algorithms have been implemented. Since our mice are inbred and of the same age, we assume that they share values of the GMRF parameters *σ*_*N*_, *σ*_*W*_, *σ*_*R*_ as well as the initial conditions *E*_0_, *R*_0_, *W*_0_, *N*_0_ and measurement errors *σ*_Pd_, *σ*_Retic_, *σ*_RBC_. Since pABA concentration affects parasite reproduction rate, we estimate a common value of *β* for each pABA treatment but allow each mouse its own inoculum size, to account for experimental variation in parasite injection volume and to allow the inclusion of data from two mice who received fewer parasites than was intended. We estimated inoculum size and *β* via multiple linear regression applied to the first 4 days of data. We estimated the remaining ten parameters (*σ*_*N*_, *σ*_*W*_, *σ*_*R*_, *σ*_Pd_, *σ*_Retic_, *σ*_RBC_, plus the initial conditions *E*_0_, *R*_0_, *W*_0_, *N*_0_) using the IF2 algorithm (Ionides et al., 2015) as implemented in the R packages pomp (King et al., 2016, 2019) and panelPomp (Bretó et al., 2019). We began by estimating the ten parameters independently for each mouse using 150 IF2 iterations of 10000 particles. Then, excluding the controls, we ran 400 iterations of the panel IF2 algorithm (Bretó et al., 2019) using 20000 particles, from each of 250 starting points distributed inside a large box in the 10-dimensional search space. We observed that these algorithms gave estimates clustered in a narrow region of parameter space relative to that spanned by the starting points. To further refine the estimates, we computed a likelihood profile over *σ*_*W*_, maximizing the likelihood over the remaining parameters at each of 100 values of that parameter. This was accomplished by starting 10 independent IF2 algorithms at each of 100 gridded *σ*_*W*_ values. Each independent IF2 consisted of 3 rounds of 100 iterations of 10000, 20000, and 40000 particles, respectively. The highest observed likelihood overall was taken to be the maximum likelihood estimate (MLE). Smoothed estimates of the state variables were obtained by running 2000 independent particle filter calculations, each using 10^5^ particles, and extracting a single trajectory from each one. Codes reproducing the entire set of calculations in detail, along with files containing the results of the intermediate calculations, will be archived permanently on datadryad.org upon acceptance of the paper.

## Acknowledgments

We thank James Fraser and the staff of Huck Institutes Flow Cytometry Facility at Penn State University, and David Adams at University of Michigan for assistance with the design and implementation of the flow cytometry protocol. We thank the members of the King, Woods, and Zaman groups at the University of Michigan for helpful comments on the manuscript. This work was supported by National Institutes of Health grants #R01AI101155 and #U54GM111274 to AAK and by a research exchange grant from to NW and AAK from the U.S. National Science Foundation-supported Infectious Disease Evolution Across Scales (IDEAS) Research Coordination Network.

## Supplementary Information

### Calculating the contribution of the different host response components to parasite fitness and host disease

We begin by making the definitions

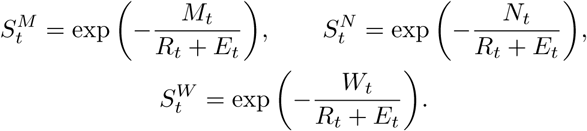

Now, to quantify the relative contribution of the indiscriminate killing response, targeted response, and parasites to RBC destruction (as shown in Figs. 3C & S-3), we calculate the following quantities that collectively sum to 1:

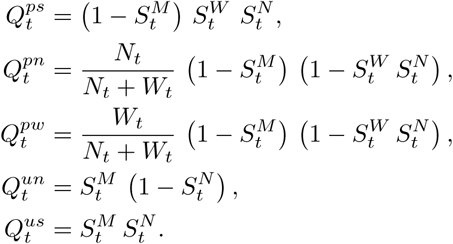

Here, 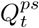 is the fraction of RBCs infected and destroyed by parasites emerging from them, 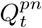 the fraction parasitized and destroyed by indiscriminate killing, 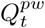 the fraction parasitized and destroyed by targeted killing, 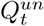 the fraction that go uninfected but are nevertheless destroyed by indiscriminate killing, and 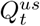 is the fraction of RBCs that survive in the preceding 24 hr.

To quantify parasite fitness, as in Figs. 3A & S-3, we divide the per-merozoite reproductive potential into

five components:

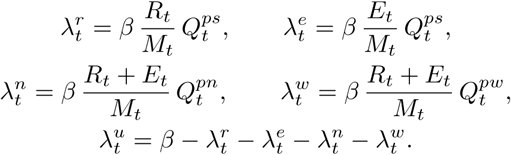

Here, 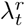 and 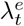 are the numbers of offspring that find and successfully reproduce within reticulocytes (immature RBCs) and erythrocytes (mature RBCs), respectively. On the other hand, we have 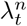, which is the unrealized potential due to destruction of parasitized cells by the indiscriminate response; 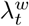, that due to destruction of parasitized cells by the targeted response; and 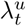, that due to lack of RBC availability. Note that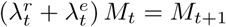 and 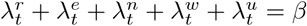.

### Calculating net change in RBC density

In general, anemia is the result of both (i) the destruction of RBCs and (ii) deficiency in their supply; if the two are balanced, then RBC concentrations remain unchanged. In Figs. 3B & S-3, we compared the previous day’s RBC losses, with reticulocyte supply on the following day, computing

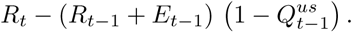

We interpret this quantity as a surplus or deficit according to whether it is positive or negative.

### Flow Cytometry

Samples were analyzed using a FlowSight Imaging Flow Cytometer (Amnis) with specifications described in table S1. 300,000 events were counted per sample and analysis performed using IDEAS software (Amnis, version 6), as follows. RBCs were distinguished from platelets and cell debris by their size (area) and CD41^-^ status. Single cells were then distinguished from doublets on the basis of their major axis (the longest dimension of the cell) and density (side-scatter). The population of single, Ter119^+^ cells was then plotted in Ter119/CD71 space and reticulocytes (CD71^+^) cells gated from erythrocytes (CD71^-^) using a gate drawn using fluorescence minus one (FMO) controls (RBCs stained with the CD41 and Ter119 antibodies but not CD71). Single color (1000 events), FMO and unstained (5000 events) controls were generated from the blood of uninfected mice and run prior to sample analysis, daily. On days 11–14 samples were not stained with the CD41 antibody, as this reagent was unavailable.

**Table S-1.** Flow Cytometer setup

| **Channel** | 1 | 2 | 3 | 11 |
| --- | --- | --- | --- | --- |
| **Laser, nm** | NA | 488 | 488 | 785 |
| **Emission Filters, nm** | NA | 430-480 | 505-560 | 560-595 |
| **Fluorochrome detected** | NA | FITC | PE | APC |

**Figure S-1.**
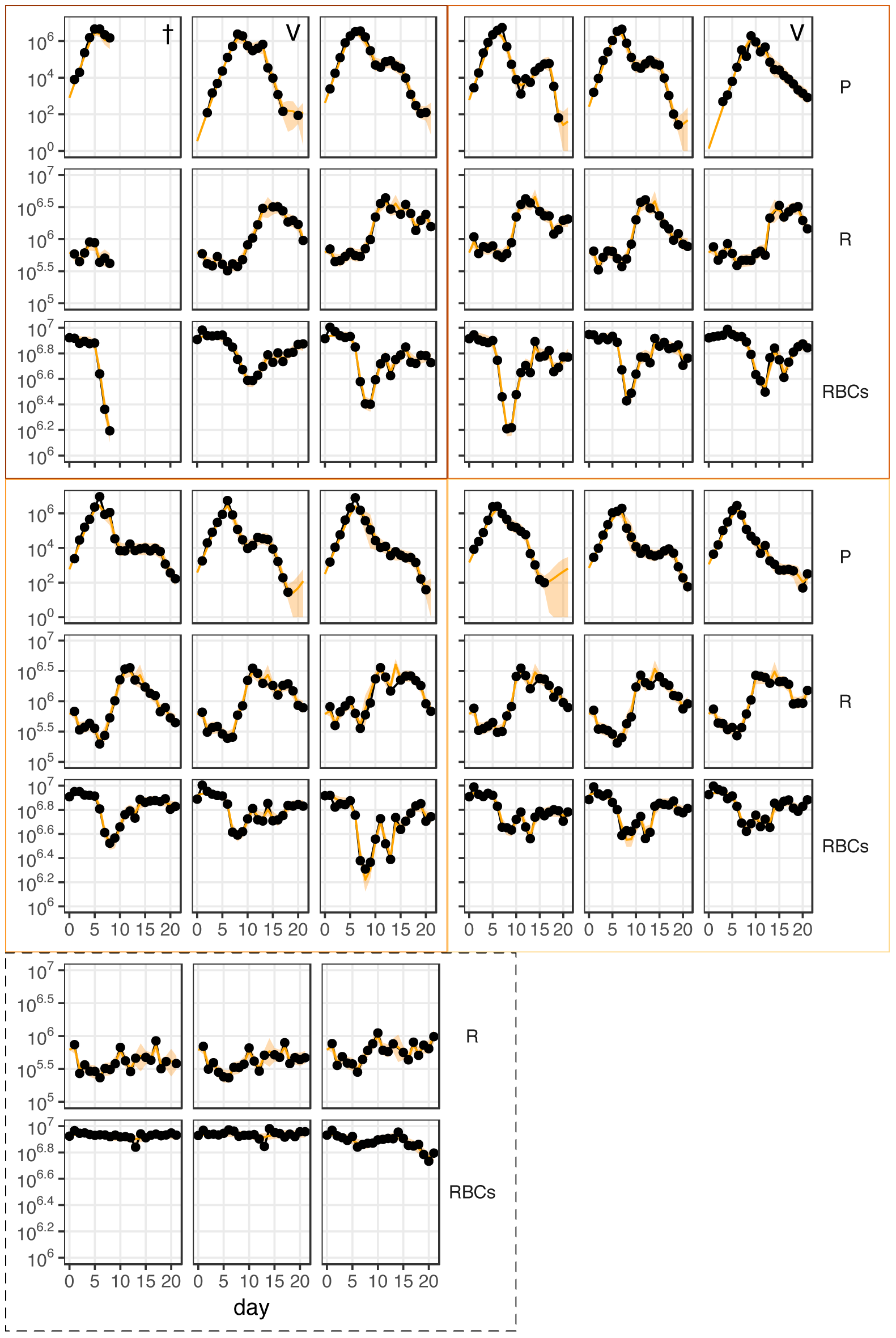
Data and fitted model trajectories of each mouse in the study. The density of parasites (P), reticulocytes (R) and total RBCs (RBCs) through time in the fifteen mice used in this study. The model (orange, smooth line) captures the data (black) well, in all cases. Twelve mice (plots with solid border) were infected with one million *Plasmodium chabaudi* parasites. These twelve infected mice were split into four groups of three that received as drinking water a 0.05% (top left block, brown border), 0.005% (top right, light brown border), 0.0005% (middle l^1^e^4^ft, orange border) or 0% (middle right, yellow border) solution of pABA, respectively. Three further mice were left uninfected (bottom left, dashed border). One mouse died (indicated by **†**) and two received fewer parasites than was intended (V).

**Figure S-2.**
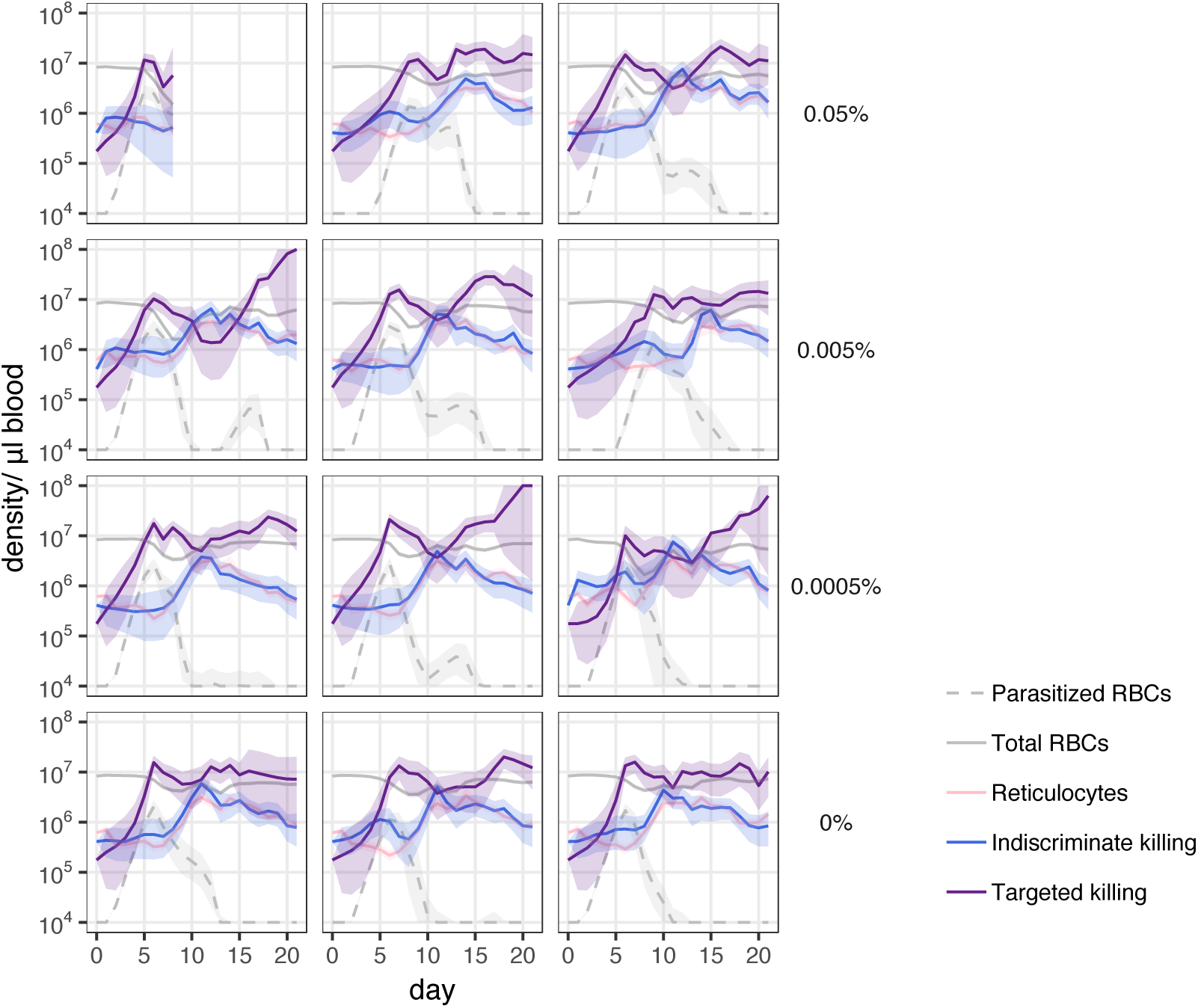
Fitted trajectories of the three components of the immune response. The trajectory of the targeted killing response (purple), indiscriminate killing response (blue) and supply response (pink) in each of 12 infected mice; the estimated densities of total RBCs (smooth line) and parasites (dashed line) are shown in grey. Plotted are the mean (solid line) and 90% confidence interval (ribbon) on the smoothed estimate of the model trajectories. Each panel shows the dynamics in a single mouse. The concentration of pABA that each mouse received as drinking water is indicated on the right. **†**indicates that the mouse died, V that the mouse received fewer parasites than was intended.

**Figure S-3.**
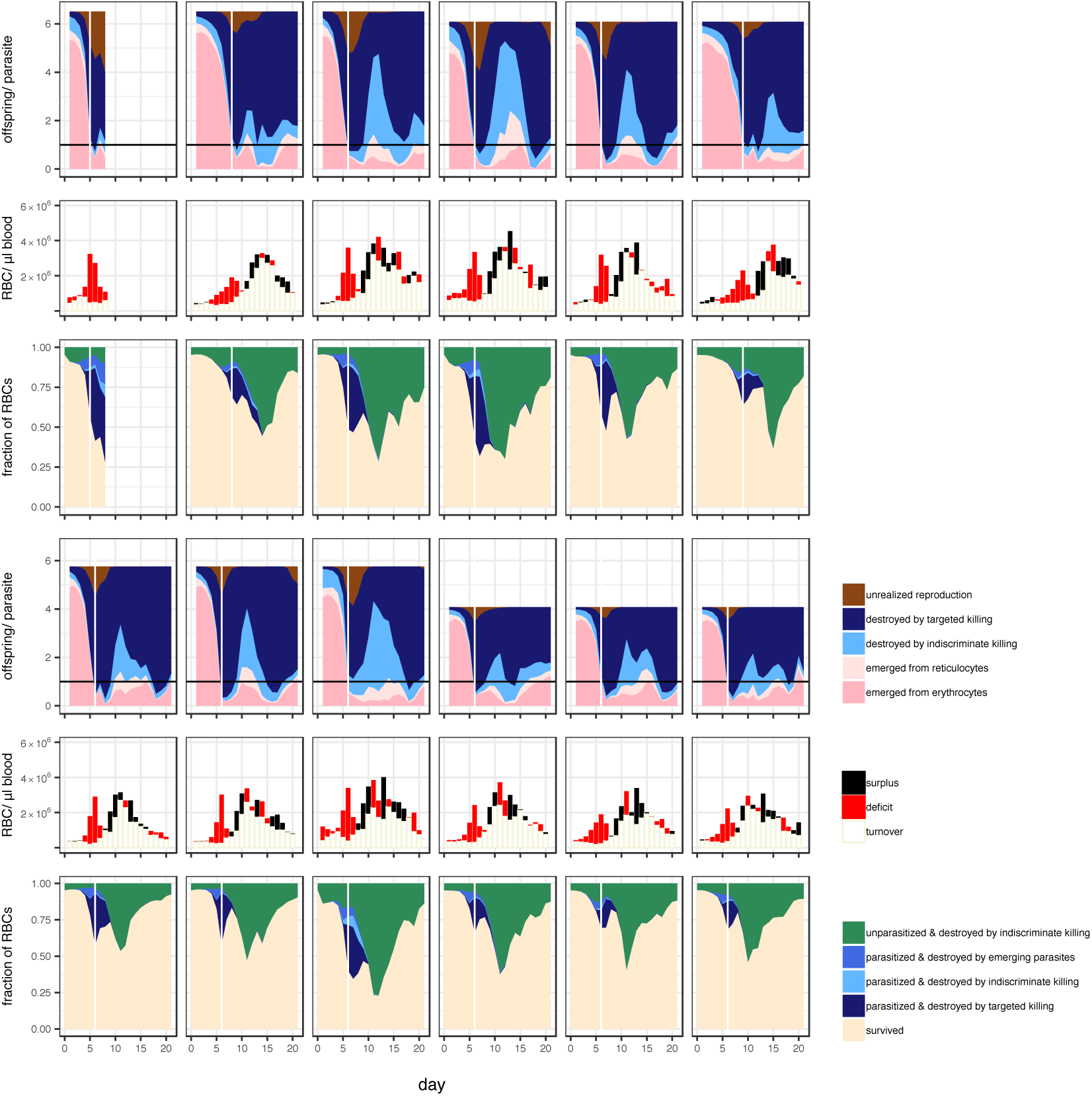
Contribution of the different components of the host response to parasite fitness and host disease in each infected mouse. Each column of three plots shows the analysis for a single mouse. First & Fourth row) Contribution of the host to (the suppression of) parasite reproduction. The height of the filled area indicates the maximum number of offspring that could be produced per parasitized cell in conditions of unlimited RBC availability. Fill color indicates the estimated number of offspring that successfully emerged (pinks), were destroyed by each of the killing responses (blues) or were not produced due to the limited availability of RBCs (brown). Black horizontal line indicates one offspring/parasite–when the blue area descends below this line the parasite population is decreasing in size. Second and Fifth Rows) Uncolored part of the bar indicates the number of the previous day’s losses that were compensated for by the present day’s supply of reticulocytes (turnover); The colored section indicates the extent to which the present day’s supply of reticulocytes exceeded yesterday’s losses (‘surplus’, black) or failed to compensate for them (‘deficit’, red). Third & Sixth Rows) The relative contribution of parasites and each of the host responses to the fate of RBCs through time. Note that parasites can contribute to RBC destruction directly, by emergence, or indirectly via the targeted killing response. White vertical line in A & C indicates the time of peak parasite density. Mice were administered a (top three rows, first three columns) 0.05%, (top three rows, last three columns) 0.005%, (bottom three rows, first three columns) 0.0005% and (bottom three rows, last three columns) 0% solution of pABA as drinking water.

**Figure S-4.**
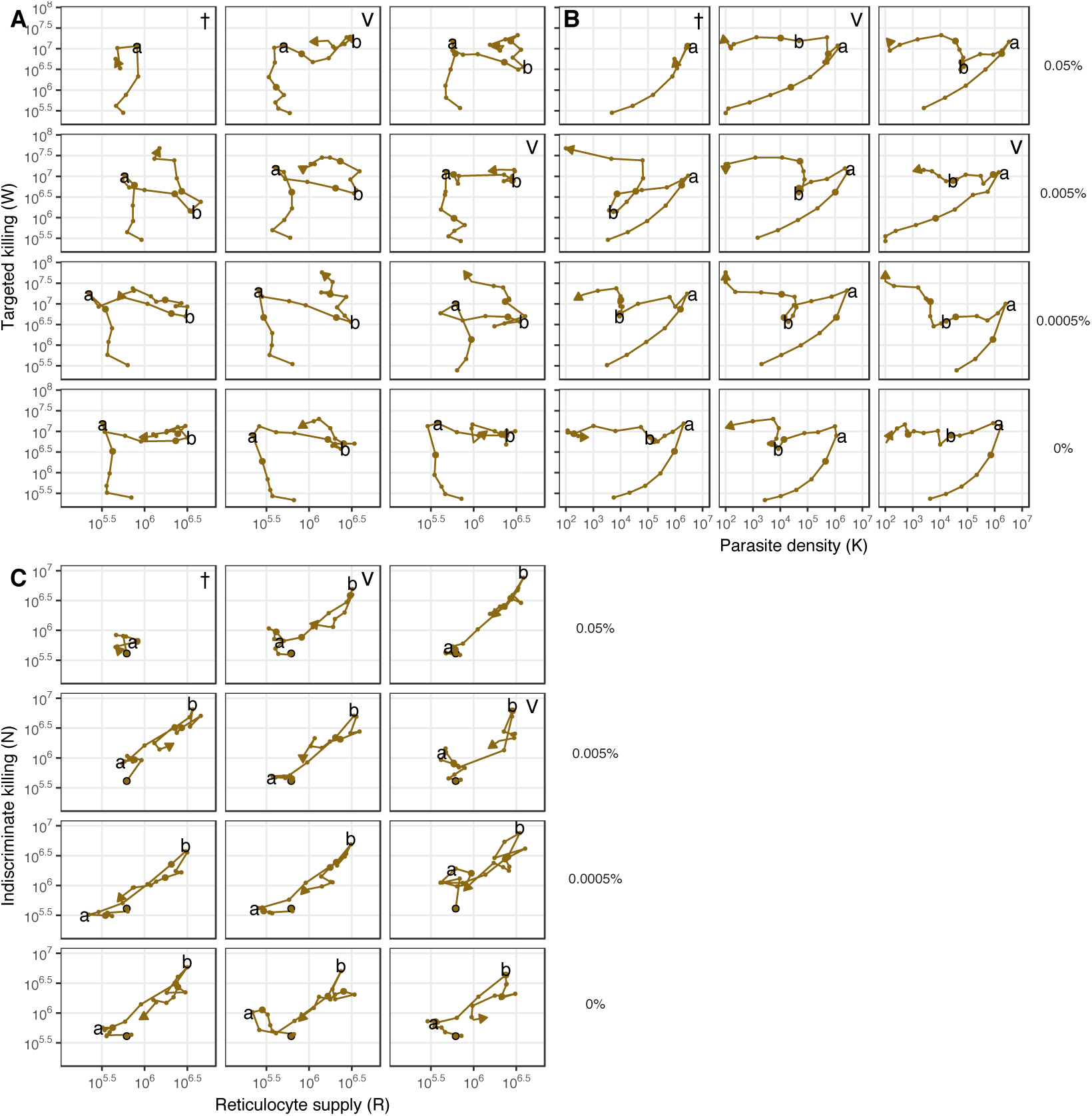
The components of the immune response are similarly deployed across mice. The relationships between A) reticulocyte density and targeted killing, B) parasite density and targeted killing and C) reticulocyte density and indiscriminate killing in infected mice given (top row) 0.05%, (second row down) 0.005%, (third row down) 0.0005% and 0% solution of pABA as drinking water. Landmarks **a** and **b** identify transitions between regimens of deployment of the components that occur. In each panel are shown the dynamics of a single mouse. The large, black-lined dot indicates the starting point of the infection; each subsequent day is marked with a dot, with large dots indicating days 5, 10, 15, 20. Densities of parasites and reticulocytes are in units per microliter of blood. Plotted are the median fitted trajectories, as estimated from the model. **†** indicates that the mouse died (and, as a result, its infection did not extend to landmark **b**), V that the mouse received fewer parasites than was intended.

